# A facile and reproducible method for the purification of peptide- and protein-functionalized DNA nanostructures

**DOI:** 10.1101/2025.06.08.658483

**Authors:** Izar Schärf, Anna Paton, Paraskevi Nani, Pol Saludes Peris, Maria Zacharopoulou, Ioanna Mela

## Abstract

DNA nanotechnology has emerged as a promising field for biomedical applications, both in the therapeutic and diagnostic domains. The ability of DNA nanostructures to carry cargos in precise numbers and orientations, makes them competitive candidates for drug delivery, biosensors or imaging agents. Two of the main challenges for translating DNA nanostructures from the laboratory to the clinic are achieving cost-effective largescale production and establishing comprehensive safety profiles. Having the ability to reliably and efficiently purify functionalized DNA nanostructures is key to both challenges, and an open question in the field of DNA nanotechnology. Here, we present a scalable method for the fast and efficient purification of high concentration of peptide- or protein-functionalized DNA nanostructures. We use a gravity-driven size exclusion chromatography approach, that has the potential to purify DNA nanostructures within 10 minutes, with yield up to 93% and purity up to 99% and is appropriate for both protein and peptide conjugates.

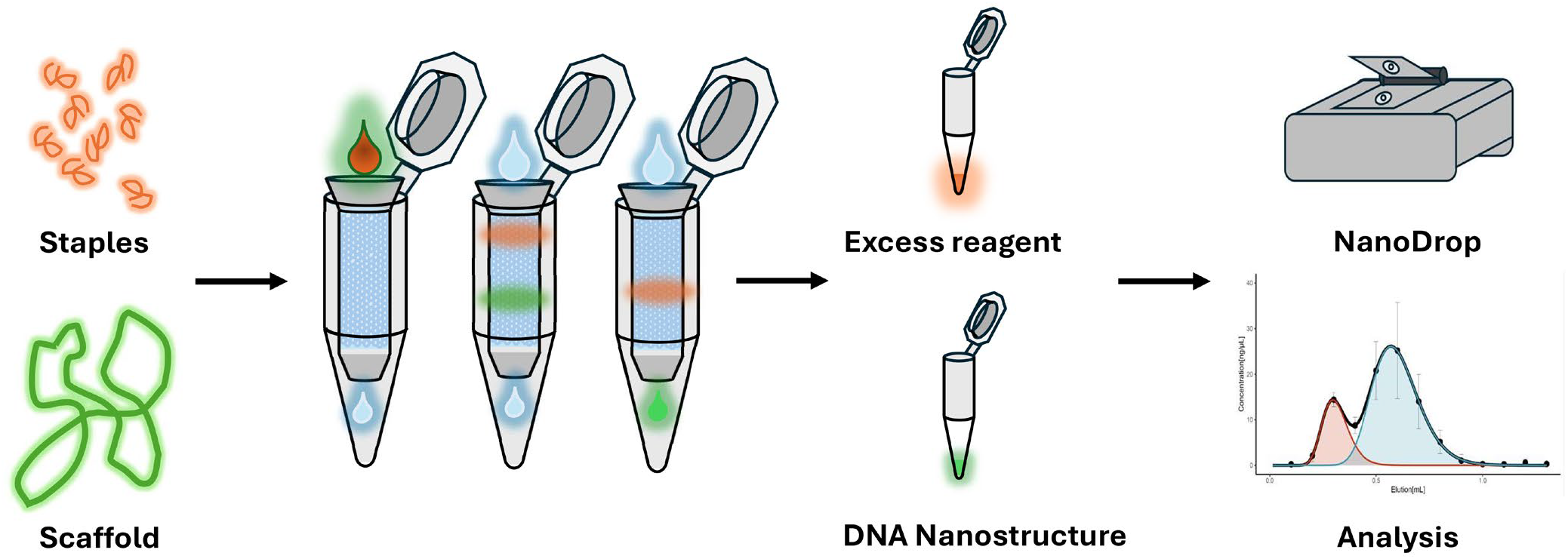

## INTRODUCTION

DNA nanotechnology offers a virtually limitless design space for creating nanoscale structures that are generating growing interest for biomedical applications, especially in the diagnostic and therapeutic delivery fields^1–5^. This can be attributed to their broad application potential, declining cost and improved design tools^4,6^. One distinct advantage of DNA nanotechnology over other types of nanoparticles, is the ability to orthogonally introduce multiple active payloads, such as proteins, peptides or small molecules on the nanostructures, while being able to precisely control their position, orientation and number of copies^7–11^. Biomedical applications of DNA nanotechnology have been well-reviewed^6,19–22^ and span a wide range, including synthetic cells, nano-motors, nanopores, bi-specifics, catalytic and enzymatic scaffolds, viral capture systems as well as drug and gene delivery systems^12– 18^.

Achieving high purity of functionalized DNA nanostructures is critical not only to ensure that they perform reliably in biomedical applications, but also to achieve regulatory approval^23^. Currently, purification of DNA nanostructures that are decorated with functional molecules (usually antibodies and peptides) in a way that is purity-, yield-, time-, and cost-efficient, can be a challenge^10,17,23–29^. Although solutions for specific applications have been explored, a universal approach to functionalized DNA nanostructure purification, similar to the widely adopted kits and workflows for DNA/RNA extraction or the robust chromatographic solutions (e.g., Ni-NTA HisTag, biotin affinity, size exclusion chromatography (SEC), or ion exchange) commonly used in protein purification, has yet to be developed^30^.

Existing methods for purifying DNA nanostructures can be inappropriate for those functionalized for biomedical applications. For example, ethanol, polyethylene glycol (PEG) or ammonium sulfate precipitation methods, are limited to DNA nanostructures with small molecule or oligonucleotide modifications and incompatible with purifying more sensitive protein-conjugated structures^24,26,28,31^. Gel electrophoresis is slow, labor-intensive, skill-dependent and non-scalable. Molecular weight cut-off (MWCO) filtration is expensive, hard to scale and can lead to structural damage and unpredictable yields, while molecular weight dialysis, though an option, is time-consuming and challenging to optimize. Both are incompatible with larger protein assemblies^25^. Methods such as Fast Protein Liquid Chromatography (FPLC) and density gradient ultracentrifugation can provide effective purification but require expensive equipment, time and high minimum working DNA concentrations making them impractical for many laboratories. Centrifugal SEC in spin columns is ineffective at removing larger proteins and might damage nanostructures sensitive to shear forces^27,32^. Gravity-driven size exclusion approaches have shown promise^29^ and especially for large-scale purifiaction^33^.

We further develop this work, to achieve small to medium scale (1–10 µg of DNA) and high concentration purification, with a gravity-driven SEC method that balances efficiency, scalability, and cost without compromising structural integrity. Concurrently, we have developed an open-access computational tool that allows for the quick quantification of purification yield and purity, based on simple NanoDrop measurements. This fills a critical gap, enabling researchers to seamlessly transition from proof-of-concept studies to larger scale experiments without changing purification systems. By addressing this gap in the range of scales, our G-SEC method bridges the needs of early-stage validation and experimental deployment towards biomedical applications.

## RESULTS

The primary focus of this work is to set up and characterize a method for quickly and efficiently purifying protein- and peptide-conjugated DNA nanostructures. We focused on 0.8 mL column volume (CV) spin columns for gravity filtration (G-SEC) using the Cytiva SuperSEC resin (Figure 1). The columns are set up in two steps, packing and equilibration (full protocol in SI). First, the column is filled with resin to achieve the target volume, in approximately 5 minutes. The resin is equilibrated with 1 CV of the elution buffer of choice for another 5 to 10 minutes. The purification is performed by adding the sample to the packed column and eluting in N fractions with desired fraction volume (Figure 1). Elution takes as little as 3 minutes and at most 15 minutes if more than 1 CV of eluent is used. The optimal loading volume (LV) for the 0.8 mL CV column was found to be 50 µL (Figure S5). To assess the scalability at higher CVs we also tested 2.5 mL CV (100 µL LV) and 5 mL CV (100 µL LV) columns (Figure S6). Testing the reusability of columns shows no deterioration in performance after 10 consecutive uses (Figure S9).

**Figure 1:**
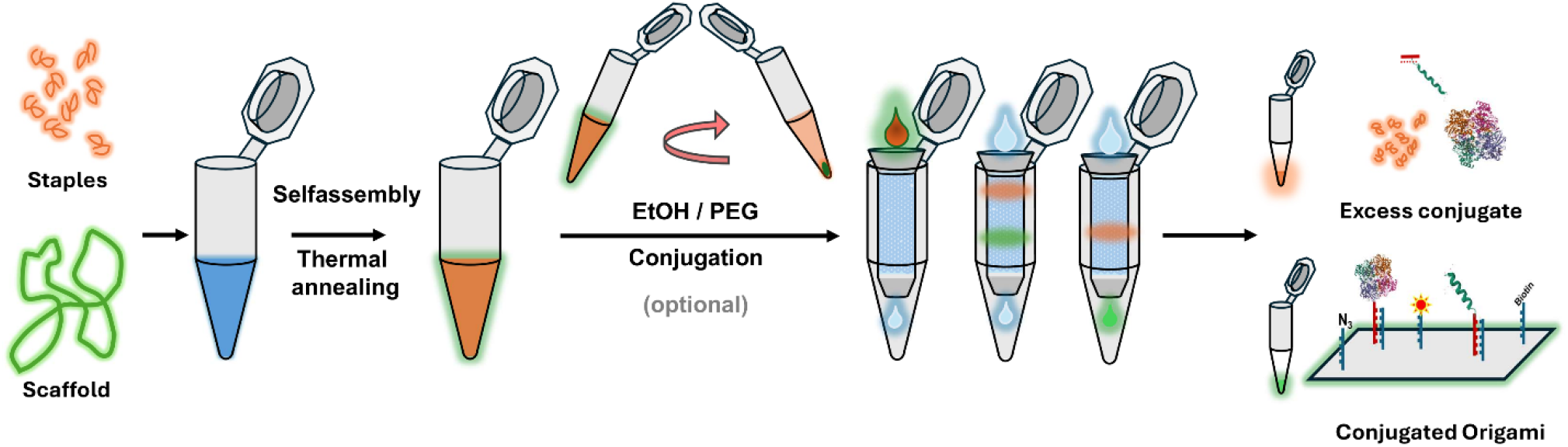
Workflow for making and purifying DNA Nanostructures. Staple ssDNA and scaffold DNA are mixed and thermally annealed, leading to the self-assembly of the nanostructures. The folded nanostructures can optionally be concentrated by precipitation by centrifuging with EtOH or PEG and then conjugated with active biomolecules of interest. The functionalized nanostructures are run through the G-SEC column which separates excess staples and conjugates from the nanostructures.

First, we set out to assess the efficiency of our method in purifying a wide range of unmodified nanostructures including different 2D (Rectangle, Triangle, 5-Well-Frame, Frame) and 3D (Tetrahedron) geometries (Figure 2). Purification efficiency was tested using agarose gel electrophoresis to assess removal of excess staples and NanoDrop measurements to estimate the amount of DNA present in each collected fraction (Figure 2.a-c). To further quantify the purification quality, we developed a simple computational approach, that allows the use of NanoDrop measurements to calculate purity and resolution of the collected fractions. These can be obtained by fitting the concentration measurements of each fraction using a sum of two log-normal distributions:

**Figure 2:**
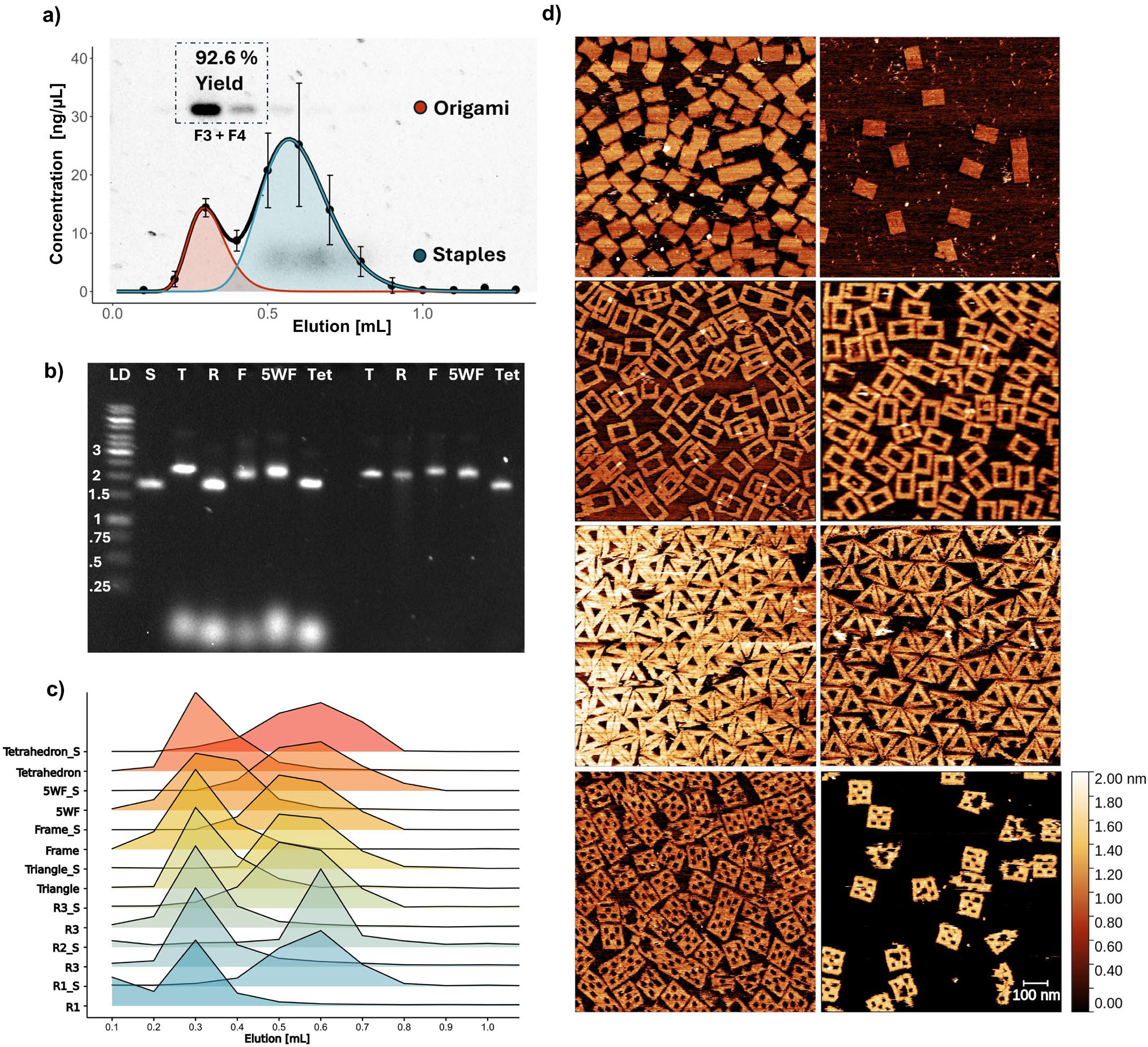
Characterization of purification efficiency on bare DNA nanostructures. **a**. Concentrations of DNA nanostructures and staple strands measured across 7 purification replicates and fitted using the sum of 2 log-normal probability density functions (Equation 1). Error bars denote the standard deviation. The sum is displayed as the black line; the two partial functions are plotted with their AUC shaded red for DNA nanostructures and blue for staples. **b**. Agarose gel electrophoresis of 5 nanostructures before and after purification (fraction 3). Scaffold (S) as control, Triangle(T), Rectangle(R), 5-Well Frame (5WF), Tetrahedron (Tet). **c**. Normalized band intensities of both DNA nanostructures and excess staples extracted from agarose gel electrophoresis images across fractions and different nanostructures. The intensities of DNA nanostructures and staples are each normalized to 1. R1 refers to repeat 1 of Rectangle till R(n), R(n)_S refers to the staple bands. **d**. Atomic Force Microscopy images of purified (left) and unpurified (right) DNA nanostructures (Rectangle, Frame, Triangle and 5-well Frame) show efficient purification and no significant damage to the nanostructures.

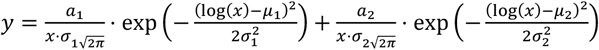

Where y is the concentration of an elution fraction as measured by NanoDrop, x is the elution volume, µ is the logarithmic mean of the distribution in x (elution volume), σ is the standard deviation of the logarithm of the elution volume and α_1_ and α_2_ are positive scaling constants.

The resolution is calculated by using the common resolution definition for close peaks, taking the full width at half maximum approach, as shown in Equation 2:

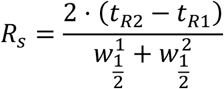

Where R_s_ is the resolution, t_Rn_ is the elution volume value at peak maximum of peak n, w^n^ = x width at half the y maximum of peak n solved numerically from the fitted partial equations from Equation 1.

Traditionally, gel electrophoresis is the only fast and reliable way of analyzing the purity and size distribution of folded DNA nanostructures. FPLC and HPLC with SEC can perform the same analysis, however, they usually have comparatively high requirements for sample concentration, which is not suitable for small scale test runs. By using the NanoDrop measurements to calculate purity, yield and resolution of purification we provide a simple computational tool for quantification of purification efficiency.

Using agarose gel electrophoresis and NanoDrop quantification, we found that elution from a 0.8mL CV column with 50 µL sample volume and 100µL fraction volume shows peak elution of nanostructures at 300µL (fraction 3) and 400µL (fraction 4) elution volume (Figure 2.a). The combined yield of nanostructures in fractions 3 and 4 measured by nanodrop is 93 ± 12% of total mass of DNA nanostructures in the sample and calculated purity of 72.2±20.2% (n=7), while the individual yield and purity in fraction 3 are 57.7 ± 6% and 98.9±1.53% respectively and for fraction 4, 35.0 ± 6.5% and 45.4±20.2%. The efficient folding of the nanostructures and separation of staple strands from DNA nanostructures is further confirmed by agarose gel electrophoresis (Figure 2b). Applying the same method to five different nanostructures (tetrahedron, 5-well frame, frame, triangle and rectangle) we can calculate the normalized band intensities of each elution fraction, separated into nanostructures and staple strands. We demonstrate the reproducibility of the method across and within nanostructures (Figure 2.c) with a triplicate purification of the rectangle nanostructure. Atomic force microscopy confirms that DNA nanostructures remain intact after purification (74.8 ± 8.4% intact nanostructures before in comparison to 76.3 ± 12.3% after, Figure 2.d).

Subsequently, we examined whether our method can be used to reliably purify peptide- and protein-conjugated DNA nanostructures that could have applications in a biomedical context. First, we tested the attachment of a tankyrase-binding peptide (TBP), conjugated to a single-stranded DNA overhang, onto DNA triangles that carry complementary DNA ‘sticky ends’. The peptide is also modified with a C-terminal Cy5 fluorophore for further detection. The DNA nanostructures were first concentrated by ethanol precipitation. We found that the recovery from ethanol purification can vary significantly yielding 79.2 ± 74% (N=6). Concentrated nanostructures were then incubated with peptides and purified using G-SEC. The efficiency of the conjugation and purification was assessed through agarose gel electrophoresis, Nanodrop measurements and atomic force microscopy (Figure 3). The yield of functionalized nanostructures was corrected for the yield of the EtOH concentration and was calculated as 64.3 ± 3.5%.

**Figure 3:**
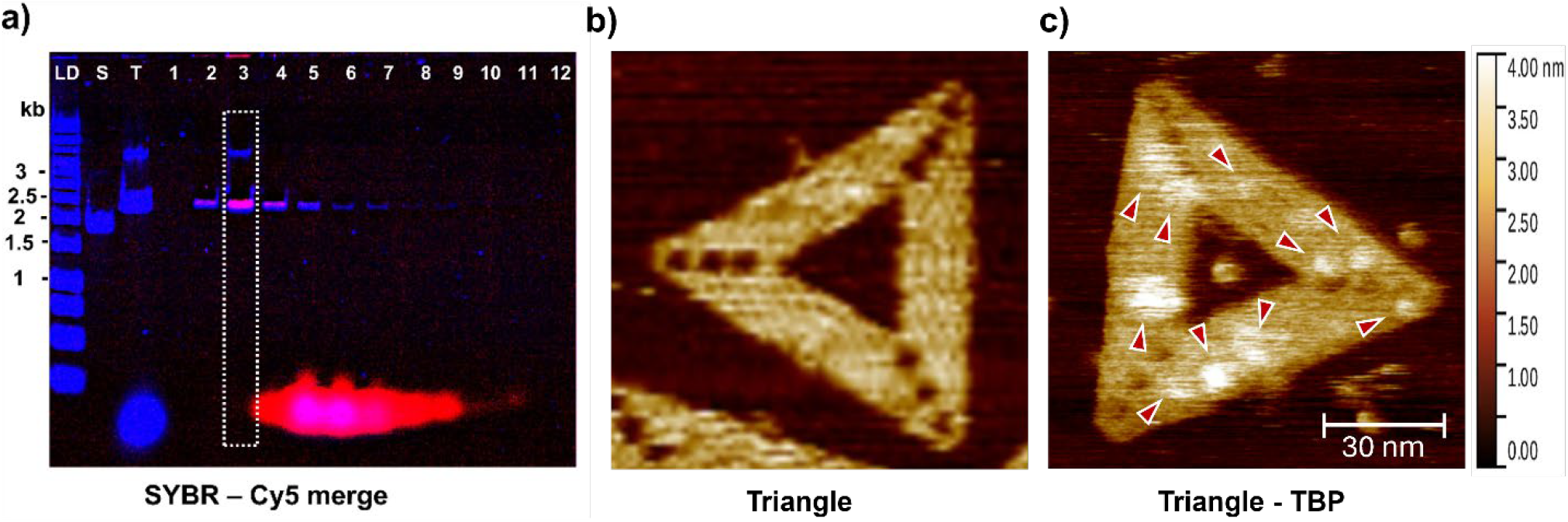
Purification of peptide-functionalized DNA nanostructures. **a**. Fluorescence image of agarose gel of peptide-functionalized DNA nanostructures. The two channels (SYBR safe (488) and Cy5 (647) show DNA and peptide respectively. Fraction 3 is highlighted by the dashed box and shows peptide-loaded nanostructures, efficiently separated from excess staples and peptides. **b**. AFM of a non-functionalized DNA triangle. **c**. AFM of a TBP-functionalized DNA nanostructure.

We then assessed the ability of G-SEC to efficiently remove higher molecular weight molecules. DNA rectangles were modified with biotinylated staples and streptavidin was used as a proof-of-principle protein. At the same time, we tested columns of different volumes (0.8mL, 2.5 mL and 5 mL) for purification efficiency in this context. For the 0.8 mL column, the recovery yield in fraction 3 was 35 ± 4% and 13 ± 2% in fraction 4, with a combined yield of 48 ± 5 %. When corrected for the ethanol concentration step, the yields are 45 ± 6% in fraction 3, 17 ± 3 % in fraction 4 and 62 ± 6% in total. Each CV shows clear separation of DNA and protein in the different fractions (Figure 4a). AFM imaging of purified samples also shows efficient purification of streptavidin-decorated DNA nanostructures.

**Figure 4:**
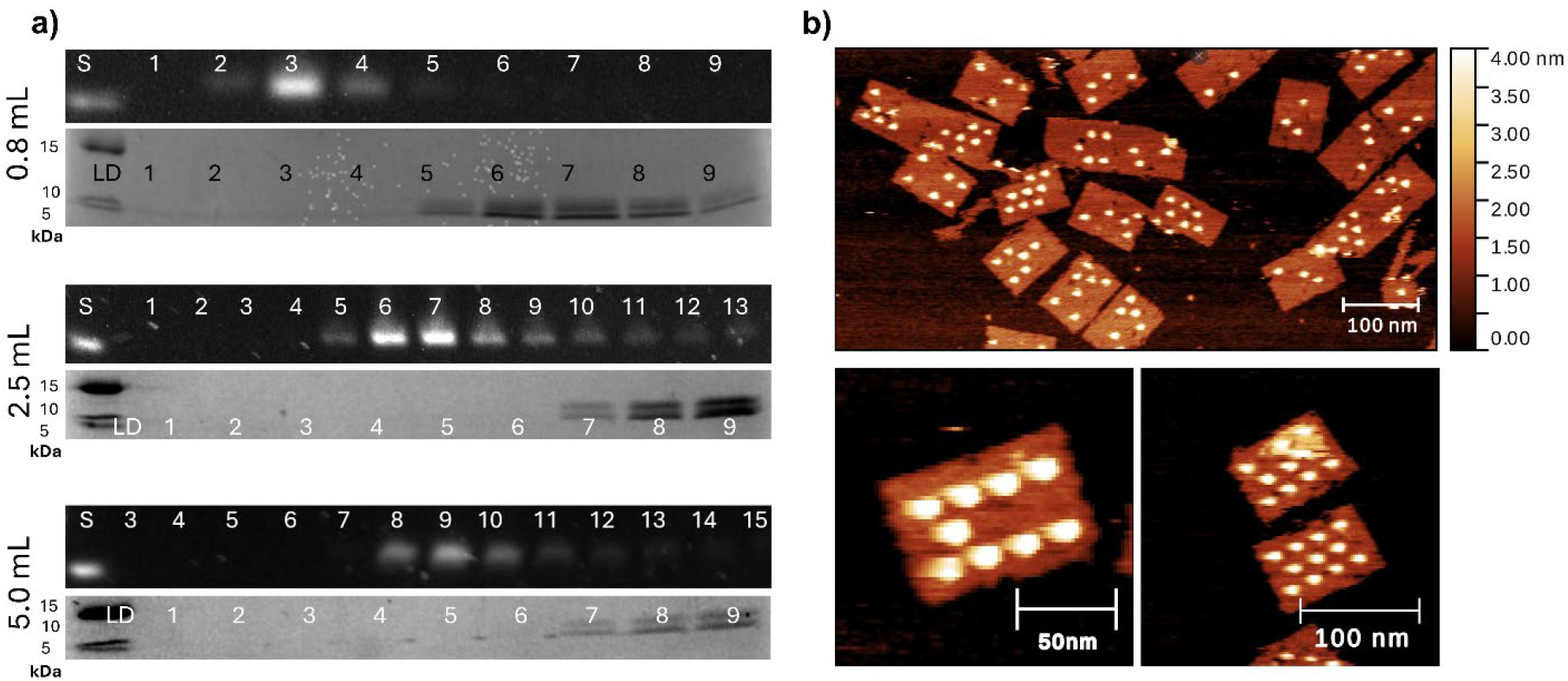
Purification of protein-functionalised DNA nanostructures. **a**. Purification comparison across three CVs (0.8mL, 2.5mL and 5mL) shows the ability to separate streptavidin from nanostructures in the aligned elutions of the DNA Nanostructures in AGE (top) and the streptavidin in SDS-PAGE (bottom). **b**. AFM of purified Rectangle-Streptavidin conjugates eluted in fraction 3 (300µL EV) on a 0.8mL CV SuperSEC column.

**Figure 5:**
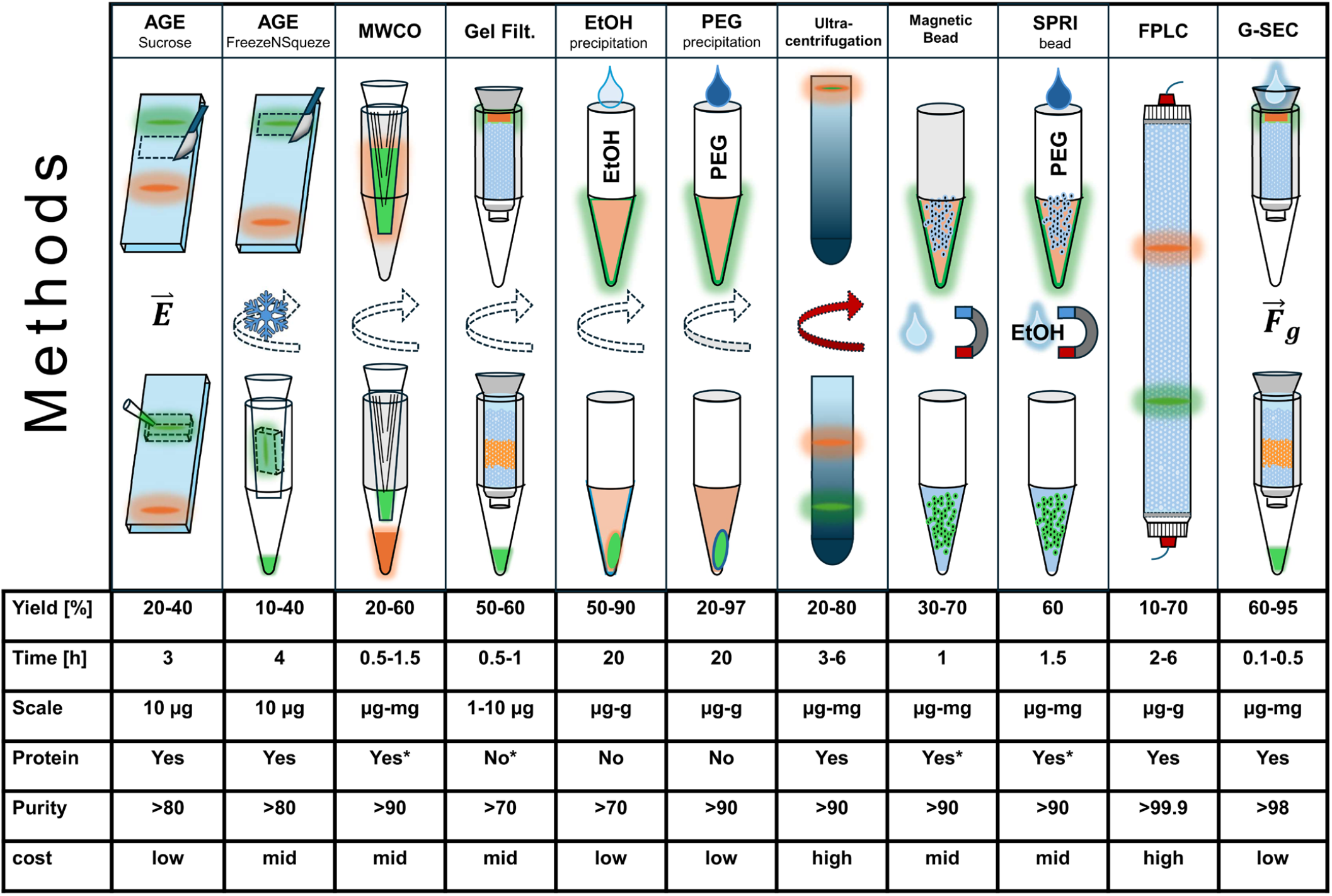
Functional schematics of most common methods for DNA nanostructures purification, comparing, the yield, time, scale, protein compatibility, and cost. Ranges are given when the literature suggests different results. * indicates that there are caveats or special cases to the method, they are however beyond the scope of this work. E indicates the electric field used to elute the sample, the snowflake indicates freezing, the turning arrows indicate centrifugation (ultracentrifugation in red) the magnets indicate use of magnets, Fg indicates gravity being the driving force

Finally, we benchmarked G-SEC against existing purification methods, using rectangular DNA nanostructures as an example. We tested extraction from agarose gels via continued electrophoresis into sucrose and ‘FreezeNSqueeze’ columns, ultrafiltration via 100 kDa MWCO filters, centrifugal gel filtration with 0.8 mL CV Sephacryl S-300, centrifugal precipitation by ethanol, FPLC-SEC with Superose 6, G-SEC with Sephacryl S-300 and Cytiva SuperSEC, (Figure S5). We compiled the results of the purification test we performed with values previously reported in the literature to provide an updated comparison of different purification methods used for DNA nanostructures.

## DISCUSSION

We have demonstrated that using 0.8 mL CV G-SEC is a fast (10 min) and efficient (up to 92.6% yield) method for purifying both bare and functionalized DNA nanostructures (yield 64%). Our G-SEC method is competitive with methods that are more expensive and time-consuming while also outperforming many in the critical categories of yield and purity. The small G-SEC approach provides a new valuable purification tool for DNA nanostructures functionalized with biologics and peptides, enabling the preparation of conjugates at concentration and purity suitable for biomedical applications. The gravity-driven nature and the washable resin make it cheap to operate, while using the Nanodrop measurements of the collected fractions for analysis of purity and resolution expedites the process further. The fact that the columns are compatible with multi-well plates, reusable and quick to elute, makes them good candidates for automated, high-throughput approaches.

## ASSOCIATED CONTENT

### Supporting Information

Methodology, Performance Comparison, AFM data, Reusability characterization, code source (PDF)

## AUTHOR INFORMATION

### Author Contributions

The manuscript was written through contributions of all authors, and all authors have given approval to the final version of the manuscript.

### Funding Sources

**M.Z**. acknowledges funding from the Ernest Oppenheimer Fund (School of Technology, University of Cambridge),

**I.M**. acknowledges funding from the Royal Society (URF/R1/221795, IES\R3\223128, IES\R2\222107, RGS\R1\231266)

### Notes

The authors declare no competing financial interest.

For the purpose of open access, the authors have applied a Creative Commons Attribution (CC BY) license.

## ACKNOWLEDGMENT

The authors acknowledge Dr Pam Rowling for her help with AKTA purification and useful discussions. Also, Cytiva, and especially David Tang for providing the Super-SEC resin.

## Notes

### Competing Interest Statement

The authors have declared no competing interest.

