## Supplementary Information for "A facile and reproducible method for the purification of peptide- and protein-functionalized DNA nanostructures"

**4 Code Availabilty** **15**

**5 Bibliography** **16**

**6 Addendum** **17**
- 6.1 Intactness counting images . . . 17

### 1 G-SEC Purification Method

#### 1.1 Purification Column Setup

The stock solution of Cytiva SuperSEC resin was thoroughly resuspended, and 25 mL were transferred into a 50 mL falcon tube, to provide a working suspension. 0.8 mL Column Volume (CV) BioRad Micro Bio-Spin columns were used for the purifications. To pack the columns, the bottom stopper was broken off, and the column was placed in a holding rack with a drip tray or an eppendorf tube, depending on column size. Care was taken for the column to be set up vertically and not to be disturbed throughout the process. 0.5 mL of resuspended resin was transferred into the column, and the solution was let to drain through. This process was repeated until the desired column volume (CV) was reached. Optionally, the column can be pre-wetted with 30-70% EtOH solution to ease the wetting of the bottom frit.

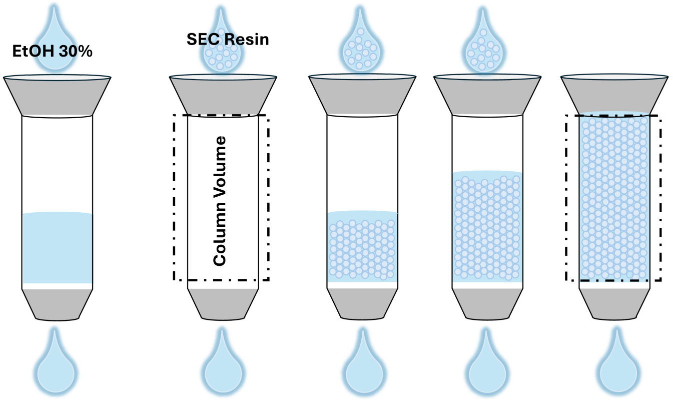

Figure S1: Preparation of a gravity SEC spin column.

#### 1.2 Purification

For purification of freshly folded DNA origami, the column was first equilibrated with 2/3 to 1 CV of elution buffer (EB). For the 0.8 mL CV columns 1.5mL EB is used for equilibration, 1 CV was similarly applied to the 2.5mL and 5mL CV columns. The buffer most commonly used is the same buffer as the one the nanostructures were folded in. If a buffer change is required, using the desired end buffer to equilibrate the column will result in buffer exchange. This sequence allowed for the fastest equilibration of the column, however if substantial buffer exchange is needed, it is advisable to use higher EB volume. All buffers used in the set-up and equilibration process were filtered through a 0.22µm filter.
Once the columns were equilibrated, the sample collection process was set-up: A series of eppendorf tubes, that would collect each purified fraction, were set up on a rack and appropriately labelled. The column was initially placed over the first tube, and the sample was gently pipetted onto the centre of the resin surface. Care was taken to ensure that no excess elution buffer was seen on the surface of the resin, the resin contained no bubbles, and that the resin had settled with a flat top, so that the purification is reproducible, and that resolution is maximized.
When all of the sample volume has run through the column, the elution process starts by adding appropriate volumes of elution fractions to the column. For the 0.8 mL CV columns, it was observed that 100 µL was the optimal volume, as lower volumes might result in slow or poor elution, as well as poor separation. Increasing the elution volume can lead to very dilute samples and worse resolution and separation. When the first fraction has been eluted, the process is repeated until the desired number of fractions is eluted. Typically, DNA nanostructures will elute within the first five fractions (0.5 CV of EB).
For the 0.8 mL CV columns, this should take 5–10 minutes, adding 1.5 mL to a 0.8 mL CV column is best done by filling the 1 mL capacity running buffer space at the top of the spin column with 1 mL EB, then 0.5 mL EB. This ensures the fastest equilibration. If the buffer change is drastic, using higher volumes for equilibration is advisable. If available, use a top frit to keep the resin flat.

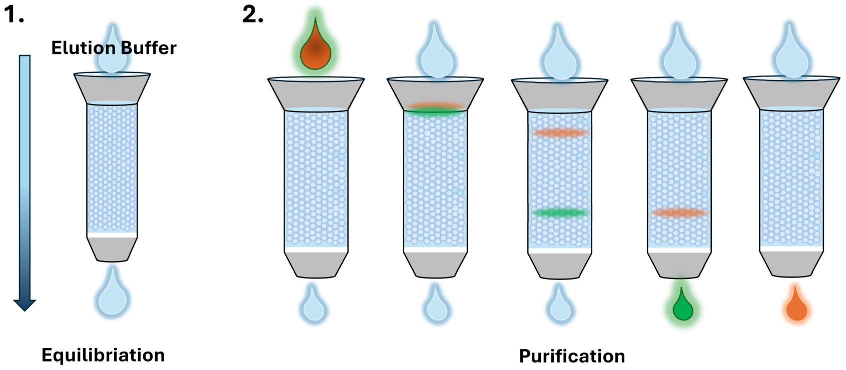

Figure S2: **Figure S2: Process of preparing a column for purification. 1.** The SEC column is equilibrated using the elution buffer. **2.** The sample is applied to the column and the sample buffer is eluted. Elution buffer is then applied in appropriate volume fractions, and the process is repeated until all desired fractions are collected.

##### 1.2.1 Required Materials

- Cytiva SuperSEC resin
    - Micro Bio-Spin™ Chromatography Columns [7326204]
    - DNA nanostructures
    - Elution Buffer
    - Eppendorf tubes
    - Pipettes 10-100µL and 0.1-1mL

#### 1.3 Analysis of Results

Analysis can be conducted by measuring the concentrations of each fraction using a spectrometer, as described in section 2.4 and fitting the results as described in section 2.10 to quantify the purity and resolution of each fraction. When the purification process is run for the first time for a new sample agarose gel electrophoresis should be used to confirm that staple strands or excess proteins and peptides are being separated from the nanostructures and that the nanostructures show a different running behavior to that of the scaffold. An unpurified sample can be run alongside purified fractions to allow for comparison of the pre- and post-purified product. The samples can then be imaged using atomic force microscopy, to assess the correct formation and stability of the DNA nanostructures.

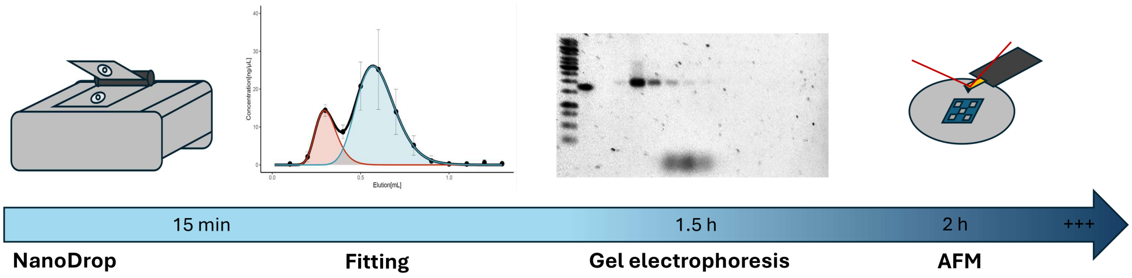

Figure S3: Standard analysis options and workflow-timeline for the purification of DNA nanostructures. Nanodrop is the fastest, taking about 1 min per sample in technical triplicates. The following analysis with an existing fitting script similarly takes a few minutes at most. Gel electrophoresis usually takes over an hour to complete. AFM can be fast although optimizing imaging can take a long time.

#### 1.4 Tips and Tricks

**Elution:** If the suspension solution does not drain one can force it to go through the frit at the bottom of the spin column by simply placing a lid on the column this should produce enough pressure to initialize the elution process. If this does not work one can try a very short spin in a slow benchtop minicentrifuge. These steps should only be done for the first addition of resin in the further steps till reaching the desired CV the resin should be left to settle and drain as vertically and stably as possible.
**Bubbles:** If a column develops bubbles in the resin, which can happen over time, especially with opening and closing cycles, the column can be re-set by vortexing the column upside-down with about 200-400µL of buffer and then turning it right-side up and letting it drain. This should reset the column for further use. This process inevitably results in losing a small amount of resin which means that the column will decrease in CV over time. Please note that this process should not be attempted during fraction elution, but only between experiments.
**Setup:** Before setting up the column, it can be washed with 30-50% EtOH solution. This will make it easier for the storage buffer of the resin to elute through the column.
**Storage:** the columns should be stored in the fridge, and if not in very regular use, they should be stored int 30% EtOH as indicated by Cytiva. The best way to store the columns and make sure they do not develop bubbles is by adding buffer the the top of the column and allowing for some buffer to start eluting. Adding the bottom cap and making sure that it is full of eluate and contains no bubbles and then adding the top cap, avoids any bubbles from being forced up through the frit when closing the bottom of the column. To open the column for use first remove the top cap then the bottom cap. For long term storage, the columns can be sealed in a 50mL Falcon tube, filled with 30%EtOH(aq) and sealed with parafilm to prevent the column from drying out.
**Cleaning:** After purifying potentially sticky proteins, the column can be cleaned using cleaning-in-place procedures. Alternatively, the resin can be collected in 50 mL falcon tubes and washed through centrifugation.
**Troubleshooting the buffers:** DNA nanostructures will run through the column differently based on the ion concentrations present in the bufffer. Using monovalent cation (e.g. Na+/ Cl-) concentration in high concentrations may disrupt unwanted interactions with the resin/proteins which might be especially relevant for some protein decorated nanostructures. DNA nanostructures have been found to interact with SEC resin and this interaction can be reduced by reducing

the concentration of Mg2+.The Mg2+ concentration can be safely reduced below 1 mM without compromising the stability of the nanostructures.[1, 2] If necessary, Mg2+ can be reintroduced after purification. The pH of the running buffer can also influence running behavior especially in protein-conjugated samples where the isoelectric point of the attached protein cannot be ignored (for example, if the proteins are naturally positively charged, they might adsorb to the highly negative DNA nanostructures).

### 2 Materials and Methods

#### 2.1 Buffers

TBE, TAE and TE were prepared according to cold spring harbour protocols and MgCl2 added to 5 mM.[3–5] DNA Origami Buffer was prepared according to Castro et al.[6]
The buffer used for protein and peptide conjugated DNA Origami purification by gravity-SEC was 150 mM NaCl 50 mM NaPO4 7 mM MgCl2, pH 7.4. All buffers were filtered either by vacuum filtaration or syringe filtration through a 0.22 m membrane.

#### 2.2 DNA Nanostrucuture folding

DNA nanostructures were folded in a MiniAmp™ Thermal Cycler (ThermoFischer Scientific) by quickly heating the sample to from 90 °C and then slowly cooling it down to 25 °C cooling with a rate of 1 C/min. Folding mix for all types of nanostructures is detailed in 2.2.

| reagent | Volume [µL] | Concentration [nM] |
|---|---|---|
| MiliQ $H_2O$ | 55 | - |
| Staple mix | 25 | 200 |
| M13mp18 ssDNA | 10 | 100 |
| Origami Buffer | 10 | 10X |

Table S3: Reagents for folding 100 µL of 10.6 nM DNA Origami

#### 2.3 Agrose gel eletrophoresis

Agarose gel electrophoresis was performed in a BioRad MiniSub Cell GT chamber at 90V for 45-90 min in an ice slurry bath. 50mL of 1% agarose gels were prepared and run with 1X TBE with 5.5 mM MgCl2 (TrizmaBase Sigma Aldrich) and 5L SYBR™ Safe DNA Gel Stain (ThermoFischer Scientific Invitrogen). DNA nanostructure sizes were confirmed with the GeneRuler 1kb DNA ladder (ThermoFisher Scientific). Gels were imaged using the BioRad Gel Doc EZ Imager, imaging condition optimization and analysis were performed using the Gel Doc Imaging Software and Image Lab (Version 6.1.0 build 7), Lane Profiles were exported with ImageJ (Version 1.54k) and plotting of ridgeline plots of intensities was performed in R.

#### 2.4 NanoDrop

DNA and protein concentrations were measured using the Implen NanoPhotometer® N60 in dsDNA mode, using the relevant sample buffer as blank. Samples were measured in technical triplicate, and the mean value is reported.

#### 2.5 Atomic Force Microscopy

Atomic force microscopy was performed on the Bruker Dimension FastScan with the FASTSCAN-D (Bruker) tips (T: 0.145µm, f0:110kHz in H2O, L: 16µm, W: 4µm, k: 0.25N/m) in fluid tapping mode. Scan parameters Tip-velocity, amplitude set-point and drive amplitude were optimized for each imaging session. For 1x1 m lines x samples /line were set to 512x512 resulting in a pixel size of 1.95 nm, 5x5 m scans were performed with 1024x1024 with a pixel size of 4.88 nm.
Images were processed in Gwyddion (version 2.63) using the following operations in this order: "Level Data by mean plane subtraction" "align rows with matching" "correct horizontal scars" "Shift minimum Data value to Zero", "false colour theme: gwyddeon.net". False colour scaling was adjusted to yield highest contrast, using "set zero to colour map minimum".

#### 2.6 SDS-PAGE

SDS-PAGE gels were prepared one day prior to use. Stacking gels were cast at 3 percent, resolving gels at 12.5 percent Acrylamide. Recipe for 2 SDS-PAGE gels as follows: Stacking gel (3%): 6.5mL H2O ,2.5 mL 4X stacking Buffer(0.5M TrizmaBase, pH6.8) , 1.0 mL Acrylamide (30%), 100µL SDS (10%), 100µL Ammonium persulfate (10%), 10µL TEMED. Resolving gel (12.5%): 3.3 mL H2O 2.5 mL 4X resolving Buffer(1.5M TrizmaBase, pH 8.8), 4.2 mL Acrylamide (30%), 100µL SDS (10%), 30µL Ammonium persulfate (10%), 10µL TEMED.

#### 2.7 ssDNA peptide conjugation, purification and hybridization on DNA Origami

Modified ssDNA (Integrated DNA Technologies, UK) carrying an azide modification at the 5' end, was mixed with an N-terminally DBCO-modified peptide (Peptide Protein Research Ltd, UK) at a 1:1 molar ratio (50 u final concentration) and incubated at 4 C overnight.
HPLC separation of the overnight azide/DBCO reaction followed (Agilent, UK). HPLC was performed using a Clarity 5uM Oligo-RP column (Phenomenex) with a 5-95 % TAE/Acetonitrile gradient. The efficiency of the overnight reaction was estimated 90 %. The oligonucleotide/peptide conjugate peak was collected, freeze-dried overnight and resuspended in DNA Origami buffer.

#### 2.8 DNA Nanotructure conjugation

The oligonucleotide/peptide conjugate was mixed with DNA origami carrying ssDNA strands in pre-designed positions complementary to the conjugate sequence (single stranded DNA overhangs) at a 10:1 conjugate:origami molar ratio. The mixture was incubated at 40 °C for 1h and subsequently cooled down to 20 C with a 1 °C/min rate.
For Streptavidin attachment, the biotinylated nanostructures were incubated for 30 min at RT with a 10X excess of Streptavidin to biotin in the sample. The nanostrucutures were first concentrated by EtOH precipitation as described in section 2.9.

#### 2.9 Purification Methods

Gravity driven size exclusion chromatography (G-SEC) was performed as described in section 1 with the 0.8 CV Micro Bio-Spin™ Chromatography Columns (7326204) and 5mL CV Pierce™ Disposable Columns (29922). The two SEC resins used were Sephacryl™ S-300 HR and Cytiva™ superSEC resin.

"Freeze 'N Squeeze" was performed according to the protocol reported in Fu, D et al 2022. Briefly, a 1% agarose gel was prepared as mentioned in 2.3. 50µL of 10 nM Rectangle, together with 10µL Loading Dye was loaded into extended loading pockets (created by excision), then run for 30 minutes. The bands of interest were excised and loaded into the BioRad Freeze 'N Squeeze™ DNA Gel Extraction Spin Columns and frozen at −20 °C as described in [7] then centrifuged to extract DNA nanostructures.

The sucrose elution method was performed according to the protocol reported in [8]. A 10mL 4% agarose gel without DNA stain was prepared, poured and left to set, then 40mL of 1% agarose gel, prepared as described in 2.3 was poured on top. An elution pocket below the bands of interest was excised, the electrophoresis tank was emptied by 20% of running buffer volume to make sure no buffer entered the excised pocket, and the gel was placed back into the chamber and the running buffer was refilled. The pocket was then filled with 30% sucrose solution containing 5mM MgCl2.

Ethanol-mediated precipitation was performed according to the protocol presented in [9]. The folded nanostrucutures were adjusted to 30% EtOH and centrifuged for 1h at 6,000 rpm. The supernatant was removed and the sample was left to dry for 10 min, ensuring that all EtOH has evaporated. The desired volume of Origami Buffer was added to the sample and it was resuspended by pipetting. The sample was incubated overnight at room temperature before use.

Ultrafiltration was performed using Amicon® Ultra Centrifugal Filters, 100 kDa MWCO. The filter was prewetted with 500 L buffer. Sample was then added, and the volume was adjusted to 450 L with origami buffer. The sample was washed three times, by centrifugation for 3 minutes at 1,000 rcf. The remaining solution was then recovered and volume measured by pipetting.

FPLC was performed isocratically with the ÄKTA pure™ using the Superose 6 5-150 increase GL SEC column. The fraction collector was set to collect 0.1 mL fractions and the flow rate was set to 0.5mL/min ensuring no overpressure. The elution buffer was 1X Origami Buffer vacuum filtered trough a 0.22µm filter. Column was equilibrated with 0.5 CV Elution Buffer.

#### 2.10 Function Fitting NanoDrop Data

Purity and resolution of the run can be calculated by fitting the peaks with 2 or more functions as shown in equation 1 as the peaks are not gaussian we use the probablility density fuction of a log normal distribution. Because of the two very different sizes (DNA Origami vs. Staples) two distributions are expected the sum of the two distributions should be able to describe the measured distributions.

$$y = a_1 \cdot \frac{1}{x \cdot \sigma_1 \cdot \sqrt{2\pi}} \cdot \exp\left(-\frac{(\log(x) - \mu_1)^2}{2\sigma_1^2}\right) + a_2 \cdot \frac{1}{x \cdot \sigma_2 \cdot \sqrt{2\pi}} \cdot \exp\left(-\frac{(\log(x) - \mu_2)^2}{2\sigma_2^2}\right) \quad (1)$$

Where y is the concentration of an elution fraction as measured by NanoDrop, x is the elution volume, µ is the logarithmic mean of the distribution in x (elution volume),  is the standard deviation of the logarithm of the elution volume and 1 and 2 are positive scaling constants.
The resolution is calculated by using the resolution definition for close peaks taking the FWHM approach as shown in equation 2. $R_s$ is the Resolution, $t_R$ is the x coordinate of the peak n, $w_{1/2}$ is the half peak maximum width.

$$R_s = \frac{2 \cdot (t_{R2} - t_{R1})}{w_{1/2}^1 + w_{1/2}^2} \quad (2)$$

We used R for fitting, plotting and calculating the purity and resolution. The results of the mean of the purification of different nanostrucutres are shown in Figure S4.
Conceptually we fit the function by visually guessing at the start parameters, altering the parameters of the individual functions till the peak height, x coordinate and width are roughly what is expected use these parameters, repeat for the second peak. If the parameters are too far off the fit will fail to find parameters; however this only needs to be done once for the initial guess of a certain CV and loading volume as the peak width and elution volume will not change much across runs and the fitting is robust enough for small changes. Purity is calculated by dividing the fraction area / fraction by total area under the curve of the fitted combined function, this is done numerically, not analytically.

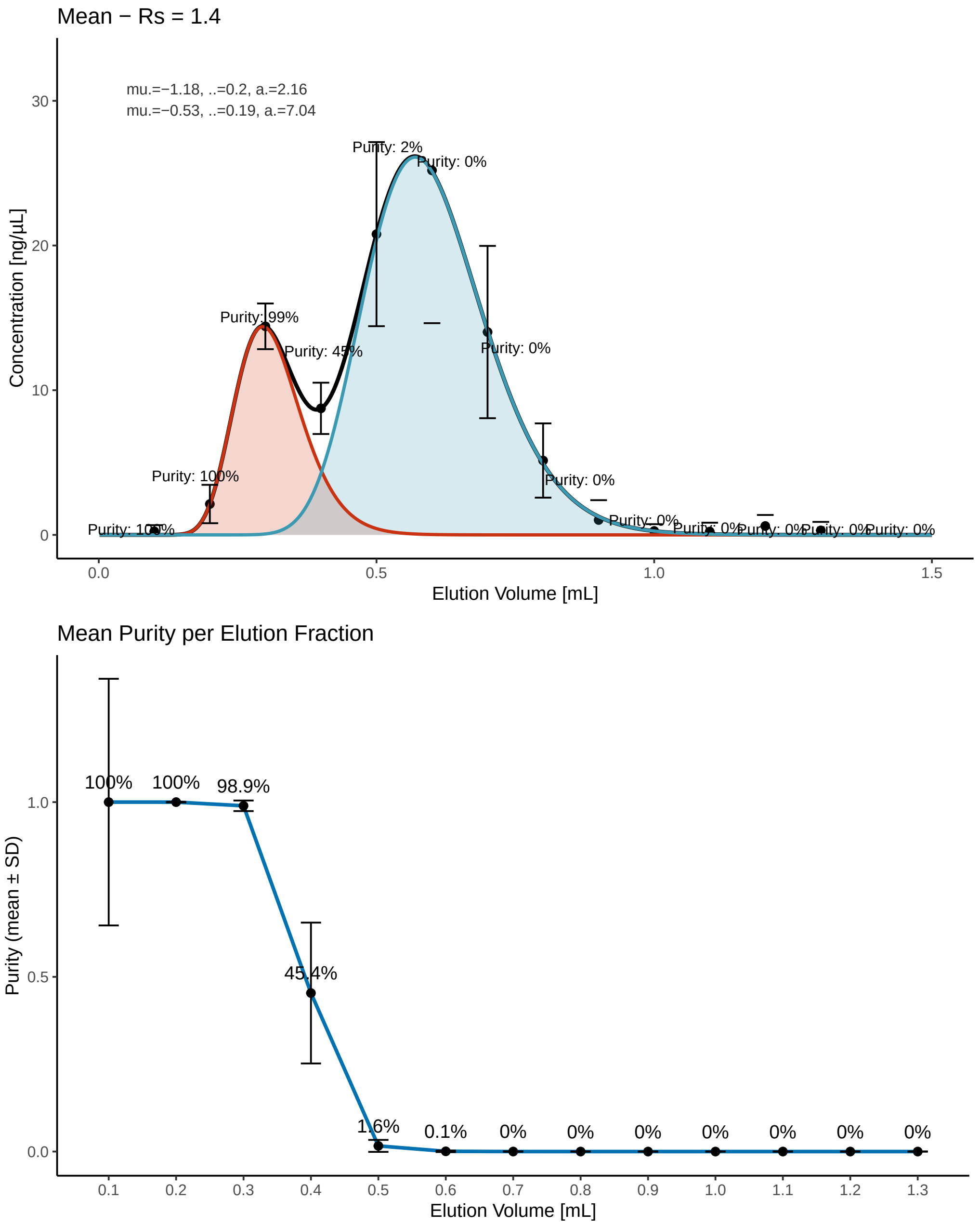

Figure S4: Plots showing the fit of the mean as well as the purity by fraction.

### 3 Additional Data

#### 3.1 Cytiva S-300 vs SuperSEC performance in 0.8 mL CV

We set out to compare the performance of purification using the Sephacryl S-300 resin to that of the SuperSEC resin in gravity driven SEC spin columns of 0.8 mL CV. We observed that staple separation from formed nanostructures is visably more efficient with SuperSEC resin than S-300 in gravity-driven SEC (Figure S5). Nanodrop concentration measurements show no clear peak separation for the S300 sample, and the agarose gel shows remaining staple strands in fraction 3 and fraction 4 (300  400 µL elution).

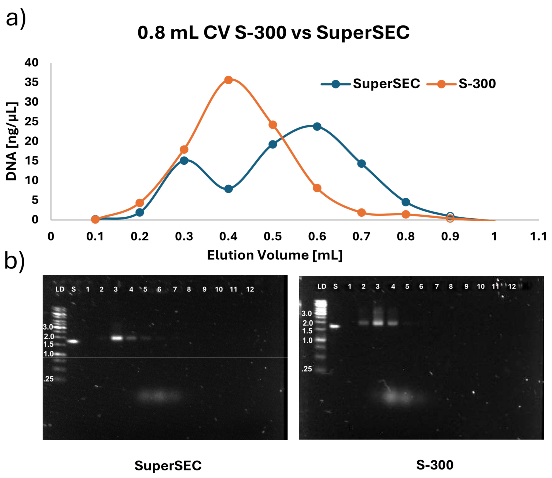

Figure S5: Comparison of S-300 vs SuperSEC size exclusion resin in Gravity-SEC applications **a)** Concentrations measured by nanodrop for 50µL 10nM (2.5 µg) R run for both resins, interpolation as visual aid. **b)** AGE images of the gels run for the samples measured in **a**on the left the gel for SuperSEC and on the right the gel for S-300.

#### 3.2 Loading volumes for 0.8 mL CV Column

We assessed the performance of the 0.8 mL CV G-SEC on 25, 50 and 100 µL loading volumes (LV). 50 µL showed the best separation at maximal loading volume. The 50µL LV delivers the best performance with highest loading capacity as shown in the comparison in Figure S6.

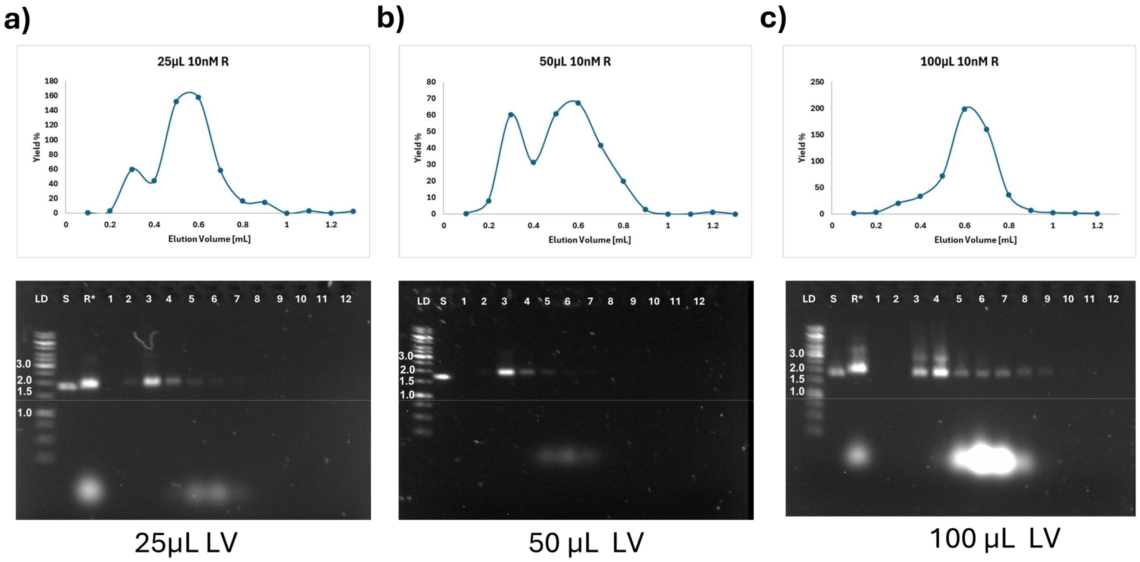

Figure S6: NanoDrop DNA concentration measurements in % yield of different sample loading volumes [a) 25µL, b) 50µL and c) 100µL] scatter plots vs the elution volume, interpolated as visual aid [upper] and representative AGE images of the fractions [lower]

#### 3.3 Column Volumes and Protein Seperation

We tested three column volumes for protein separation using 10 nM of Rectangle nanostructures: 0.8 mL (50µL loading volume, 100µL fractions), 2.5 mL (100µL Loading Volume, 125µL fractions) and 5mL (100µL loading volume, 250µL fractions). The results of the comparison are shown in Figure S7. Examples of the columns are shown in Figure S7a Separation and elution peak breadth increases with increasing CV as seen in Figure S7b while dilution increases with rising CV. Streptavidin is easily separated from both the Rectangle and the Rectangle-streptavidin conjugates as shown in Figure S7c for the example of 0.8 mL CV and similarly higher CVs separate the Nanostructures from the Streptavidin as shown in Figure S7d. From this we conclude that using higher CVs than 0.8 mL is contra productive for small volume purifications of about 50 µL although higher loading volumes such as shown in Figure S6c might result in lacking separation.

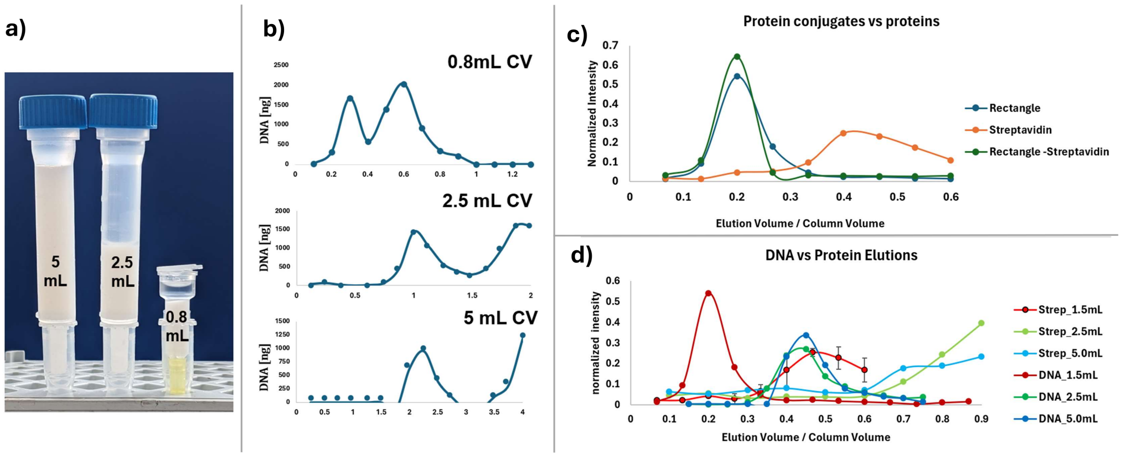

Figure S7: **a)** shows a picture comparing 5 mL, 2.5 mL and 1.5mL CV spin columns on 1.5mL eppendorf tubes. **b)** Shows the total DNA weight estimated by nanodrop concentration measurements for purifications from the 5 mL, 2.5 mL and 0.8 mL CV versus the elution volume. **c)** Shows the normalized intensity extracted from gel images of agarose gel electrophoresis and SDS-PAGE of the Rectangle Nanostructure (AGE), the Rectangle conjugated to Streptavidin (AGE) and streptavidin (SDS-PAGE) versus the elution volume divided by the column volume. **d)** Shows the normalized intensity of DNA Nanostructures in AGE and Streptavidin in SDS-PAGE vs the Elution Volume divided by the column volume.

Complimentary to Figure S7 we show the comparison of NanoDrop concentration measuremets for the fractions of the unconjugated, Streptavidin conjugated rectangle and the TBP conjugated Triangle below in Figure S8. Notably both the Triangle and Rectangle were EtOH precpitated before conjugation, 500 µL 10.6 nM 25 µg Nanostrucutre + 217µL EtOH, resuspended in 100 µL Origami buffer. 30 µL of the resuspended nanostructures were then conjugated with streptavidin/TBP and purified on the 0.8 mL CV SuperSEC columns with PBS +7 mM MgCl2 as described in section 2.1. The difference between the Rectangle and Triangle in the main elution peak is minimal, yet the nanodrop clearly shows the ssDNA-TPB being seperated from the Triangle nanostrucutre, this was also demonstrated in main paper Figure 3a).

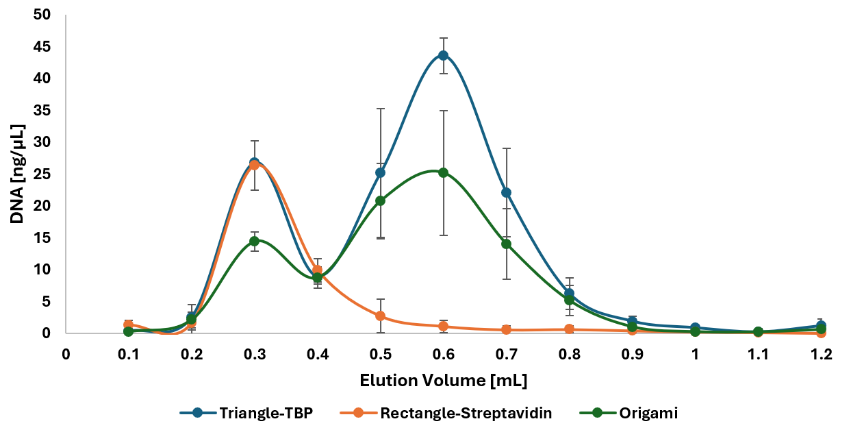

Figure S8: Showing a comparison of the concentration measurements of Triangle-TBP conjugate (N=3), Rectangle-Streptavidin conjugate (N=3) and the mean (N=7) of all nanostructures as shown individually in S10 gravity purified on the 0.8 mL CV SuperSEC column.

#### 3.4 Nanostrucuture integrety

To assess the effect of purification on the structural integrity of the nanostructures, we counted the intact vs non-intact structures before and after purification. This was performed with 5WF, frame, triangle, and rectangle. The annotated images to the data in S8 are shown in the addendum. Note that the AFM tip can destroy nanostructures which means that areas scanned for longer will have increasing numbers of defective nanostructures. The counts were performed with clickmaster2000 : https://www.thregr.org/wavexx/software/clickmaster2000/

| Nanostructure | Intact | Broken | Intact (%) | Broken (%) |
|---|---|---|---|---|
| 5WF | 1212 | 753 | 61 | 38 |
| 5WF F3 | 311 | 245 | 56 | 44 |
| Frame | 61 | 13 | 82 | 17 |
| Frame F3 | 47 | 11 | 81 | 18 |
| Triangle | 75 | 17 | 81 | 18 |
| Triangle F3 | 65 | 17 | 79 | 20 |
| Rectangle | 1937 | 612 | 75 | 24 |
| Rectangle F3 | 754 | 84 | 89 | 10 |

Table S8: Showing the counts and percentages of Nanostructures divided between intact and defective.

#### 3.5 Purification of different Nanostructure

As previously shown in main paper Fig 2 d) we tested the 0.8 mL CV SuperSEC columns for several nanostrucures, the unpurified vs purified strucutres are shown with their corresponding gels below in Figure S9. The concentration measurements of the samples run on the gels in Figure S9 are shown in Figure S10.

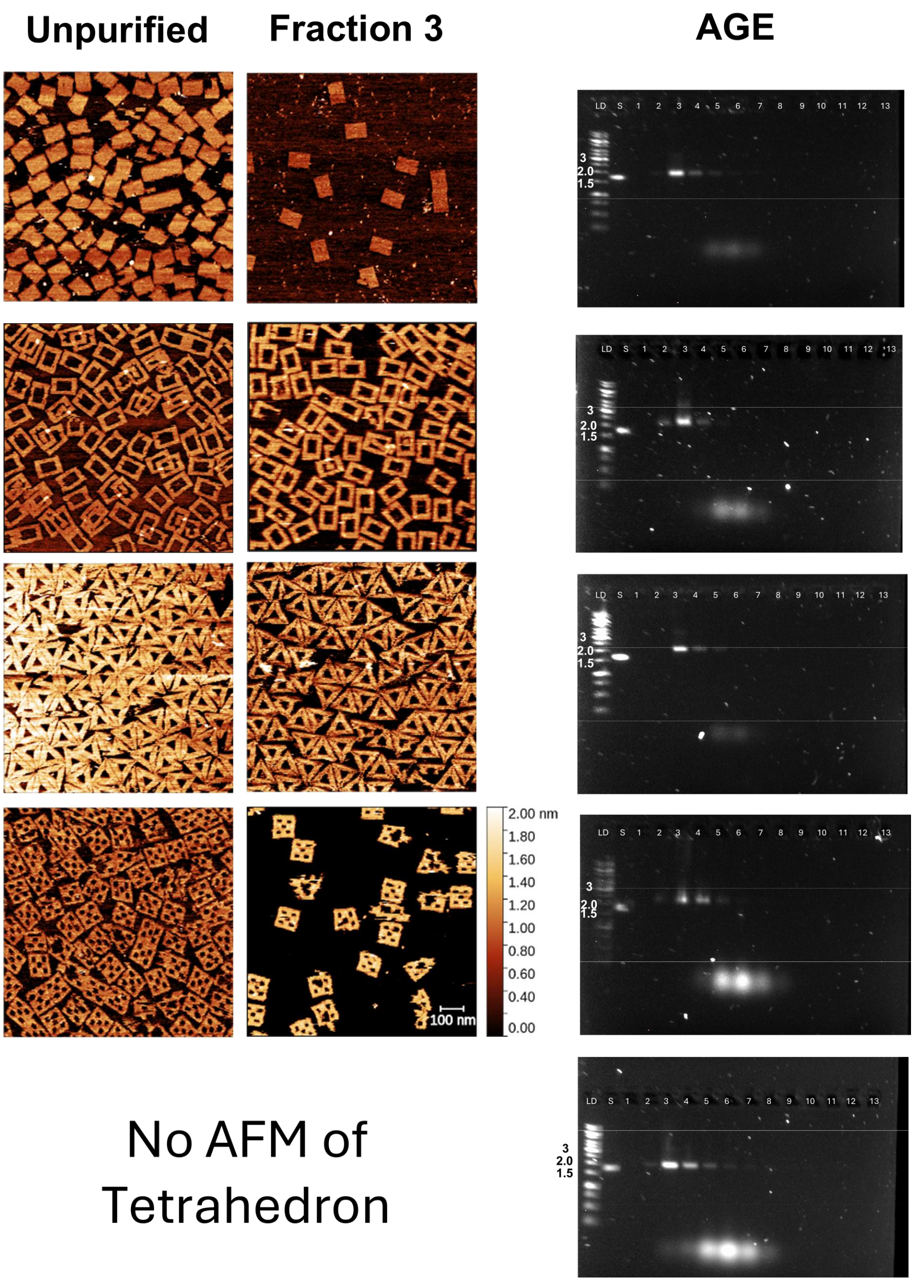

Figure S9: Showing the AFM of unpurified vs purified nanostructures for rectangle, frame, triangle, 5-well frame and tetrahedron* and the gels for their respective purifications. 50µL of 10nM folded nanostructure were purified by a 1.5 mL CV SuperSEC column with 100µL elution fractions. *The AFM of the tetrahedron structure is missing as the structure's geometry and lack of large surface area for mica interaction does not lend itself to AFM imaging.

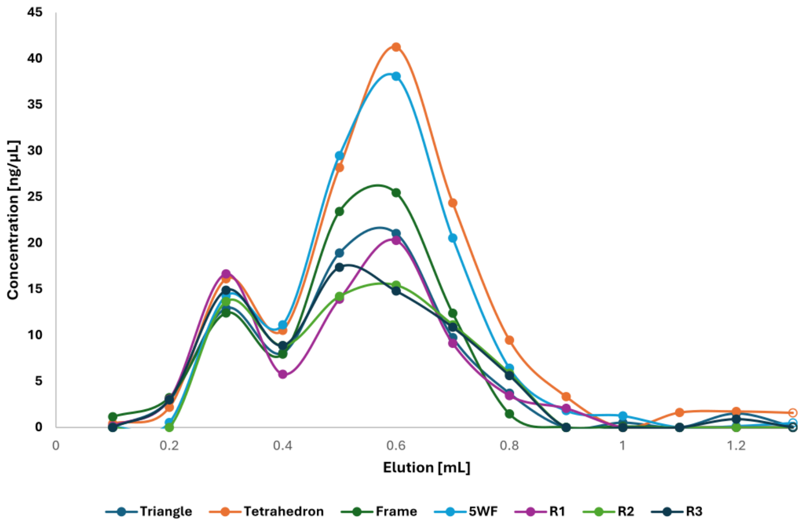

Figure S10: Shows the concentration measurements vs elution volume for 50 µL 10nM LV on the 0.8 mL CV SuperSEC with 100µL fractions for each of the Nanostructures shown in Figure S9. Three replicates are included for the Rectangle nanostructure as R1, R2 and R3. The interpolation serves as a visual guide only.

#### 3.6 Reusability

To test the reusability of the columns by clean in place methods common to SEC columns for FLPC systems, we washed the 0.8 mL CV SuperSEC columns with 1 CV elution buffer after each purification of DNA Nanostructures. We used columns up to 10 times before returning the SEC resin to a falcon tube for collecting the resin from several columns to be washed more thoroughly by NaOH and stored in 30% EtOH for reuse. The problem that forced the retirement of all set up columns was the formation of air bubbles in the column. While it is possible to reconstitute a column by vortexing it so the resin resettles we chose to retire columns instead. Below in Figure S11 we show the first (Rectangle) and tenth (Triangle) purification of DNA Origami on a 0.8 mL CV column.

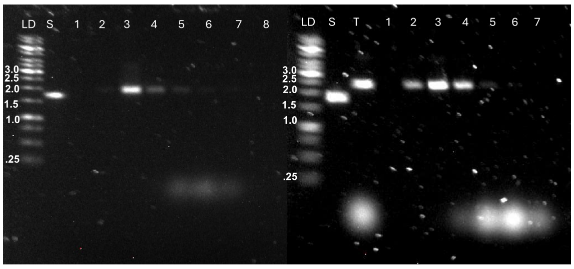

Figure S11: Reusability: For a 1.5mL CV SuperSEC column first (left - Rectangle) and tenth (right -Triangle) purification are shown.

#### 3.7 Purification Method Comparison

We tested several methods for purifying DNA Origami, the two common Gel extraction techniques: Sucrose Elution and FreezeNsqueeze, precipitation via Ethanol and the Ultrafiltration by Molecular weight cut off (MWCO). The methods were performed as described in section 2.9.
The Sucrose elution method can work quite well for functionalized nanostructures S12a however the method is highly user dependent and difficult to repeat, new users will struggle getting it right the first few times. Its even harder when one does not have a punch to standardize the cut out process for the elution pocket. Over 7 repeats the sucrose elution method yielded 38+-22% of input.
Freeze and squeeze S12a also relies on cutting out the minimum amount to extract the whole gel segment containing the desired nanostructure. This method recovered 38+-47% on input nanostructures. In several cases no recovery was possible making this method unreliable at best.
Both of these gel extraction methods are clearly not scalable and both contaminate the DNA

irreversibly with a DNA stain aswell as presenting low recovery and significant reliabilty issue. MWCO (100kDa) filtering had varying results yielding 65+-16%. In our earlier tests loading 25 µL 10nM 5WF we did not recover any DNA Origami, in tests with Streptavidin we saw no separation as the Streptavidin still interfered with AFM imaging.
The purification by Ethanol was the method with the highest variability, from recovering almost nothing S12b EtOH(1) to recovering 200% S12b EtOH(4) final mean yield of 79+-74% of 2.5µg Origami in 50µL (10nM), all yields are uncorrected for purity. Recovering DNA Origami from anything less than a 50µL (2.5µg ) starting volume / mass is very hard as there is seldomly a visable pellet. The method provides very varying degrees of purity as can be seen in S12b as well as having an often unreliable recovery. However the reliability increases with scale, using the method on 500µL Rectangle DNA Origami recovered a pellet in every case. This is why we reccomend this method as a concentration step rather than a purification step, with higher amounts of starting material and a tolerance for impurities.

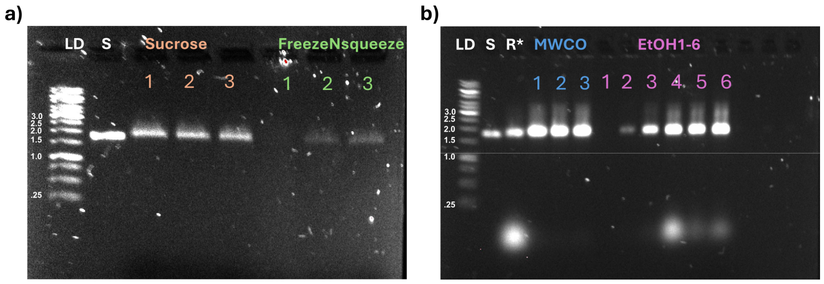

Figure S12: Shows four alternative methods we tested. a) shows the Eluting into Sucrose from an AGE method and the FreezeNsqueeze method both in triplicate. b) shows the Molecular weight cut of filtration by centrifguation in triplicate as wellas a sextuplicate of the precipitation by EtOH. Scaffold DNA is marked by “S” and unpurified Rectangle is marked by “R*”.

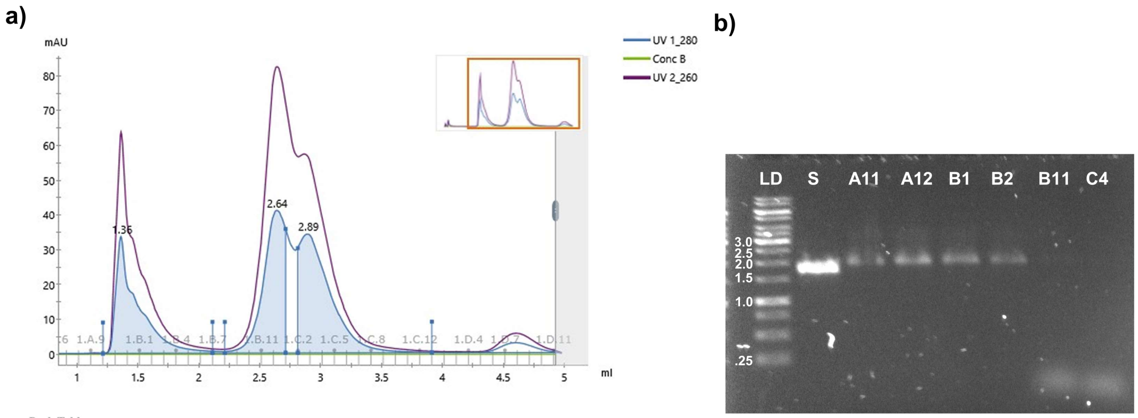

Peak Table

| Peak | Retention (ml) | Area (ml*mAU) | Area % | Ext coeff. (mg ⁻¹ ml cm ⁻¹) | Fraction(s) | Volume (ml) |
|---|---|---|---|---|---|---|
| Peak A | 1.358 | 6.714 | 27.18 | | 1.A.10 - 1.B.6 | 0.900 |
| Peak B | 2.637 | 7.972 | 32.27 | | 1.B.8 - 1.B.12 | 0.500 |
| Peak C | 2.892 | 10.02 | 40.55 | | 1.C.2 - 1.C.12 | 1.100 |

Figure S13: Shows a representative purification run of 100µL 10 nM freshly folded Rectangle purified on the Äkta-Pure with a 5x150 Superose 6 SEC column, the Origami containing fractions represented by peak A yield 36% with fraction A12 at 1.1mL elution volume containing the highest concentration with 4.5 ng/µL represeting a 12X dilution at 9% recovery. b) An AGE image of the first peak from A11 to B2 and the two following peaks B11 and C4 are shown in comparison to the scaffold, 20µL is loaded per fraction, 5 µL Ladder and 5µL 10nM ssDNA scaffold.

### 4 Code Availabilty

Code is available at the following link: https://github.com/izarscharf/FittingNanoDrop/

### 6 Addendum

#### 6.1 Intactness counting images

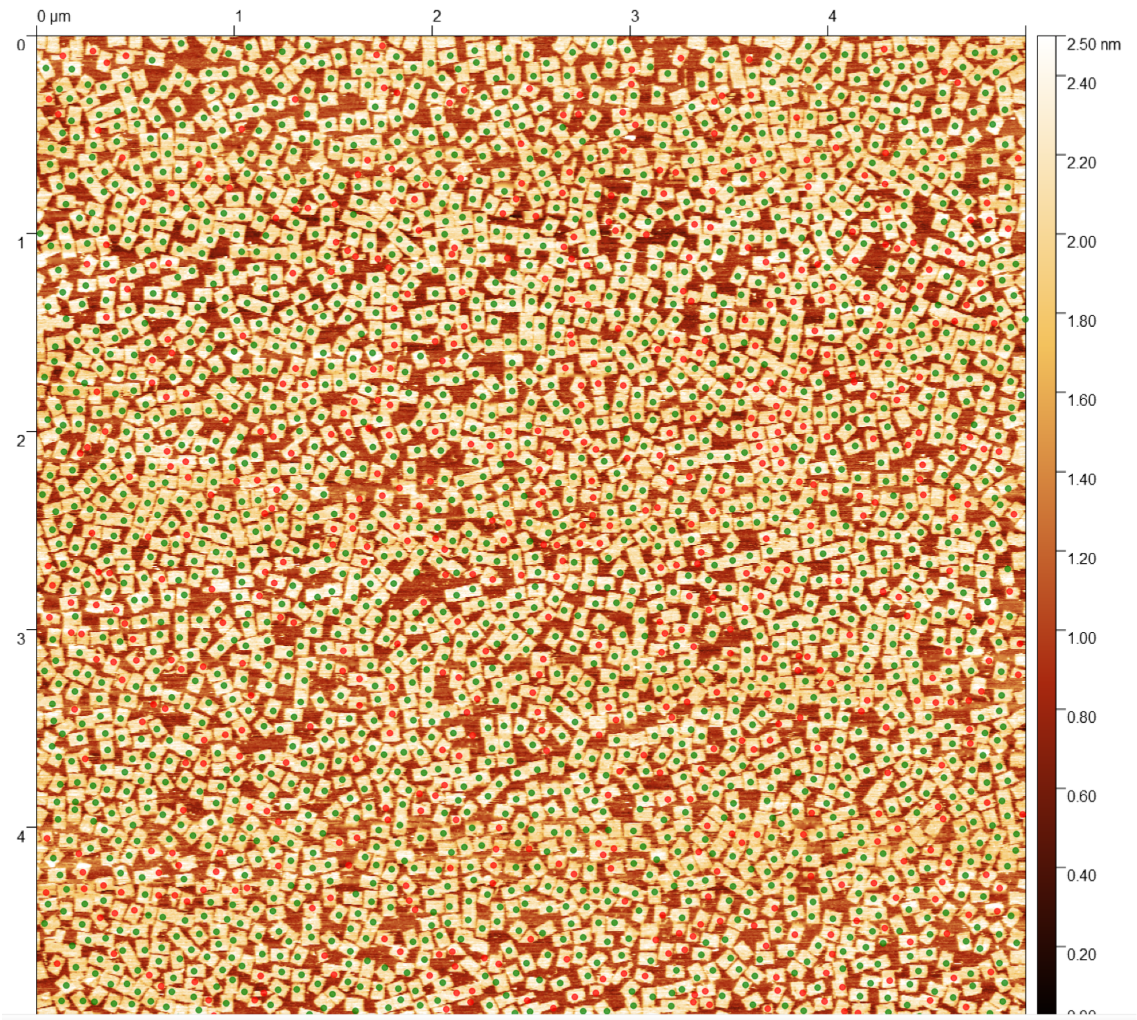

Figure S14: Rectangle unpurified 5x5µm AFM image

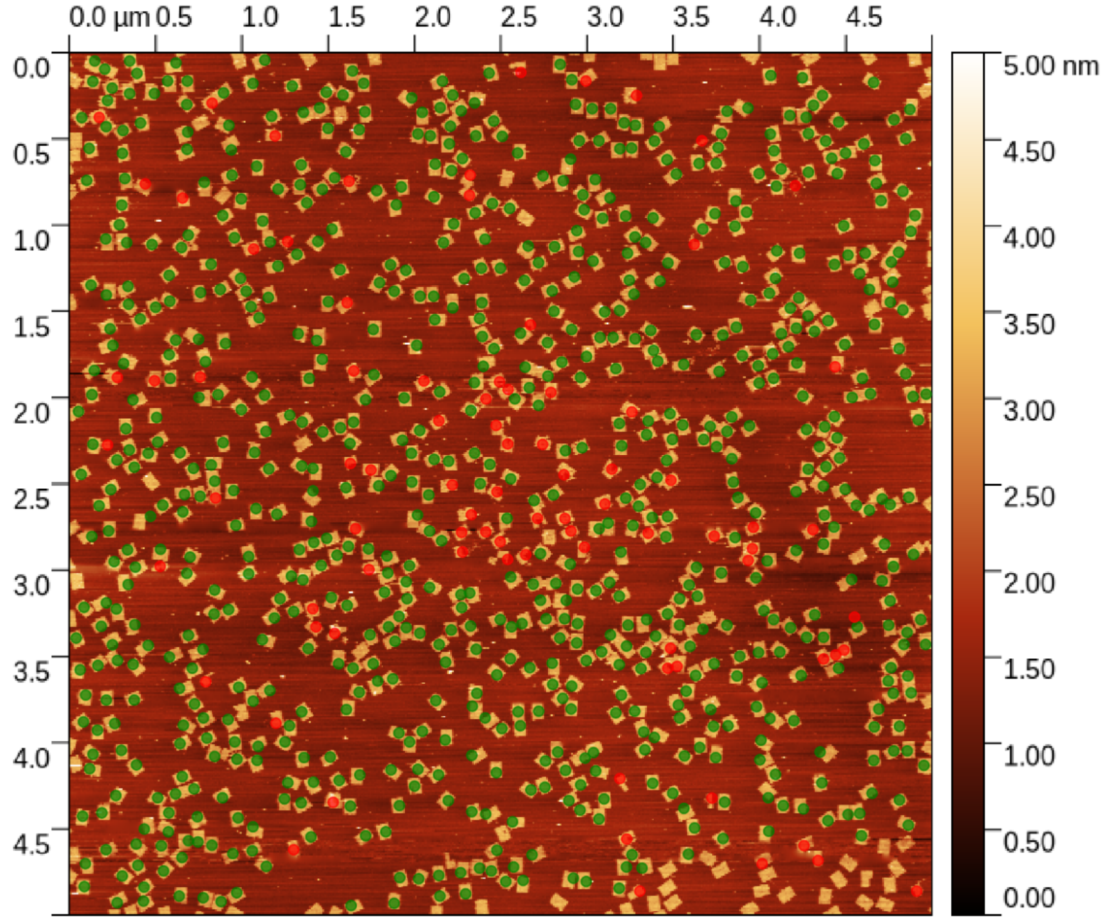

Figure S15: Rectangle purified Fraction 3 (300µL EV)5x5µm AFM image

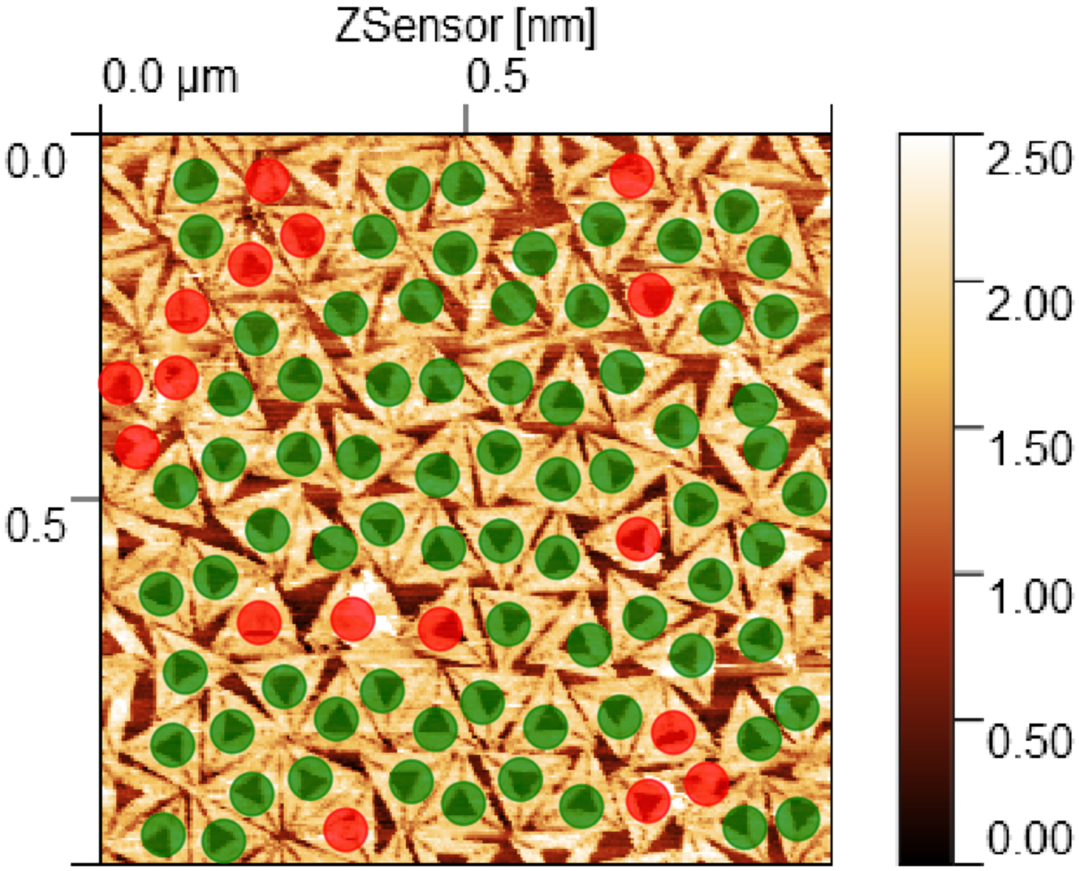

Figure S16: Triangle unpurified 5x5µm AFM image

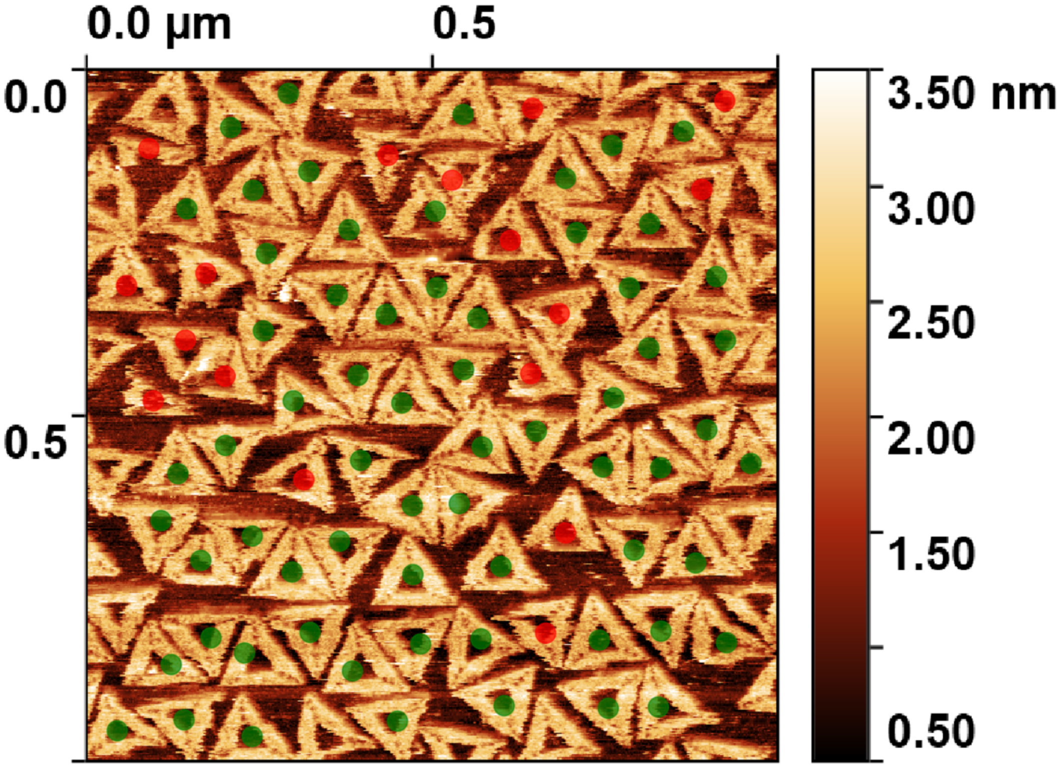

Figure S17: Triangle purified Fraction 3 (300µL EV)5x5µm AFM image

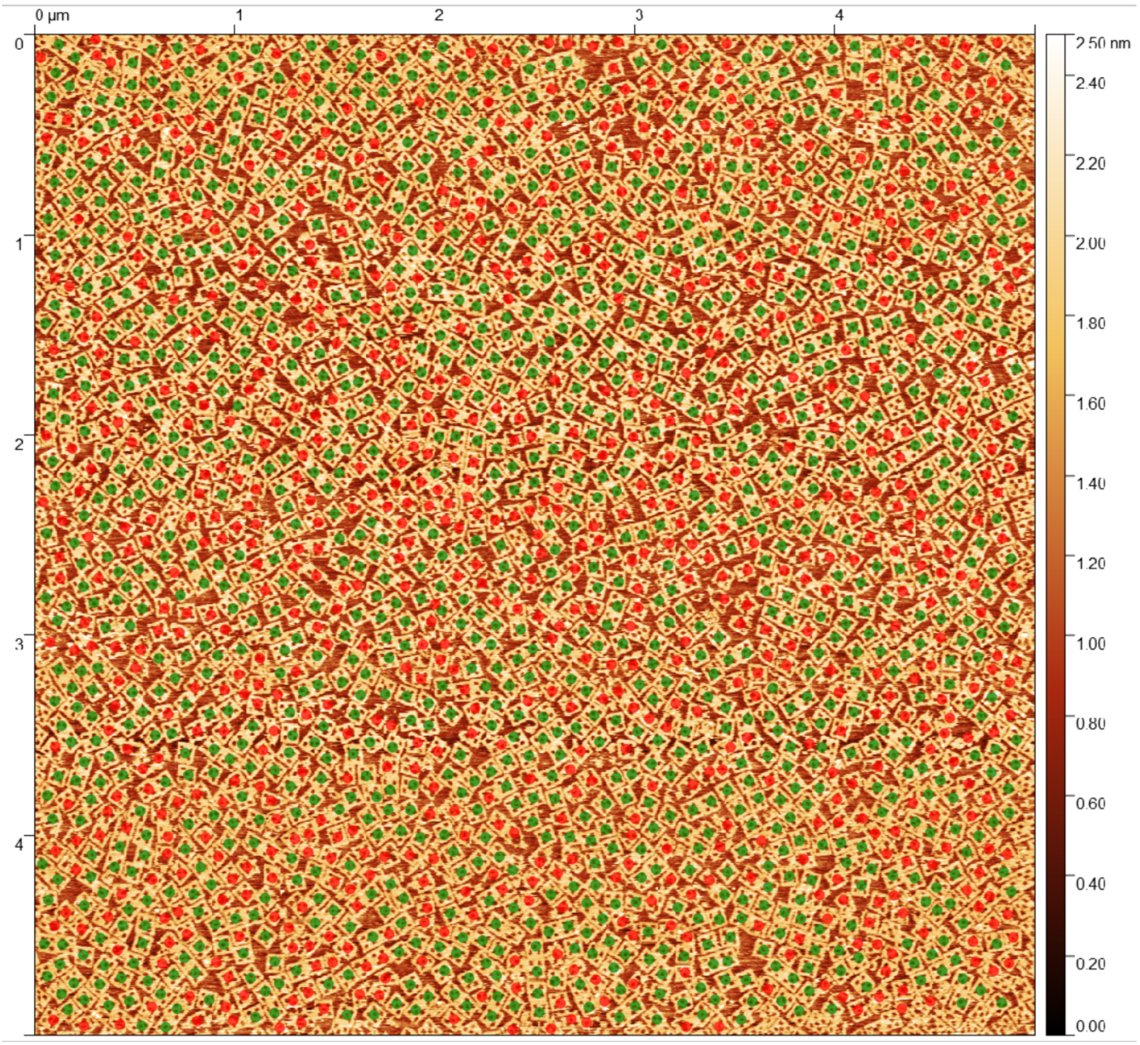

Figure S18: 5-well Frame unpurified 5x5µm AFM image

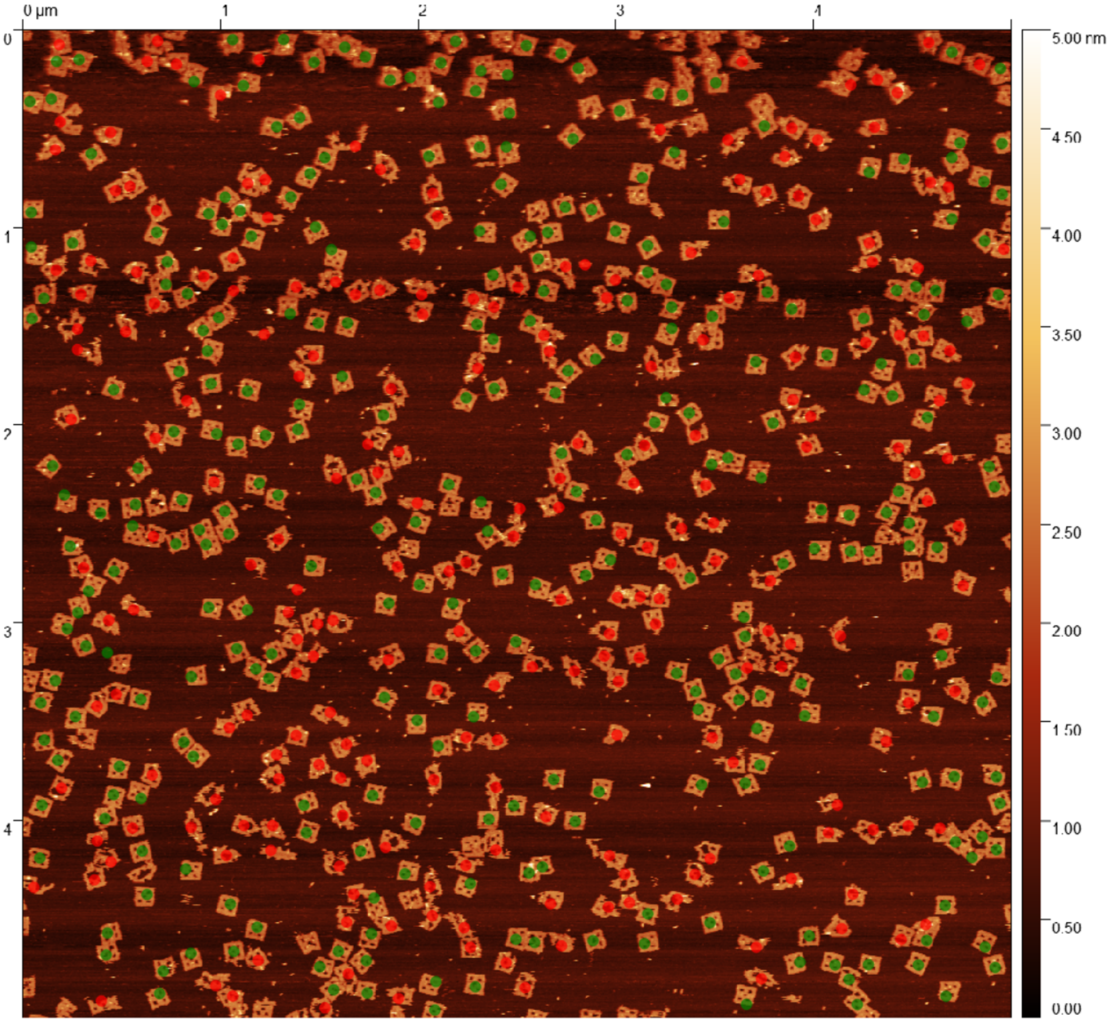

Figure S19: 5-well Frame purified Fraction 3 (300µL EV)5x5µm AFM image

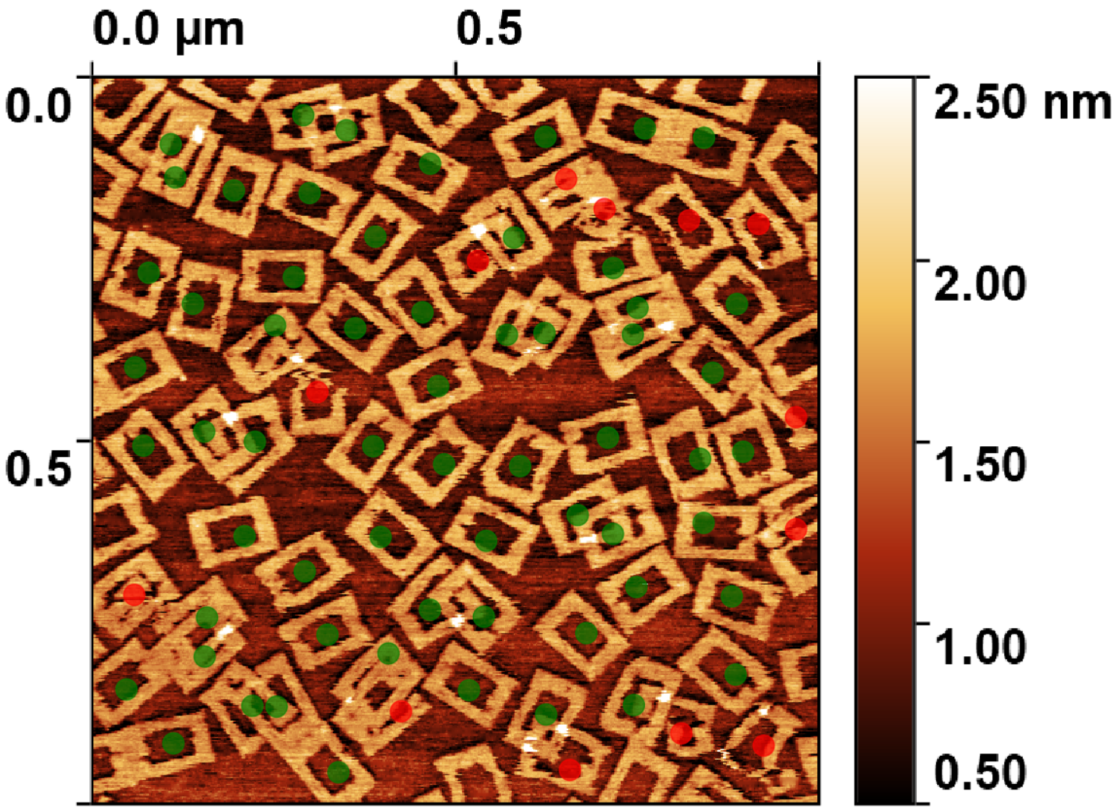

Figure S20: Frame unpurified 5x5µm AFM image

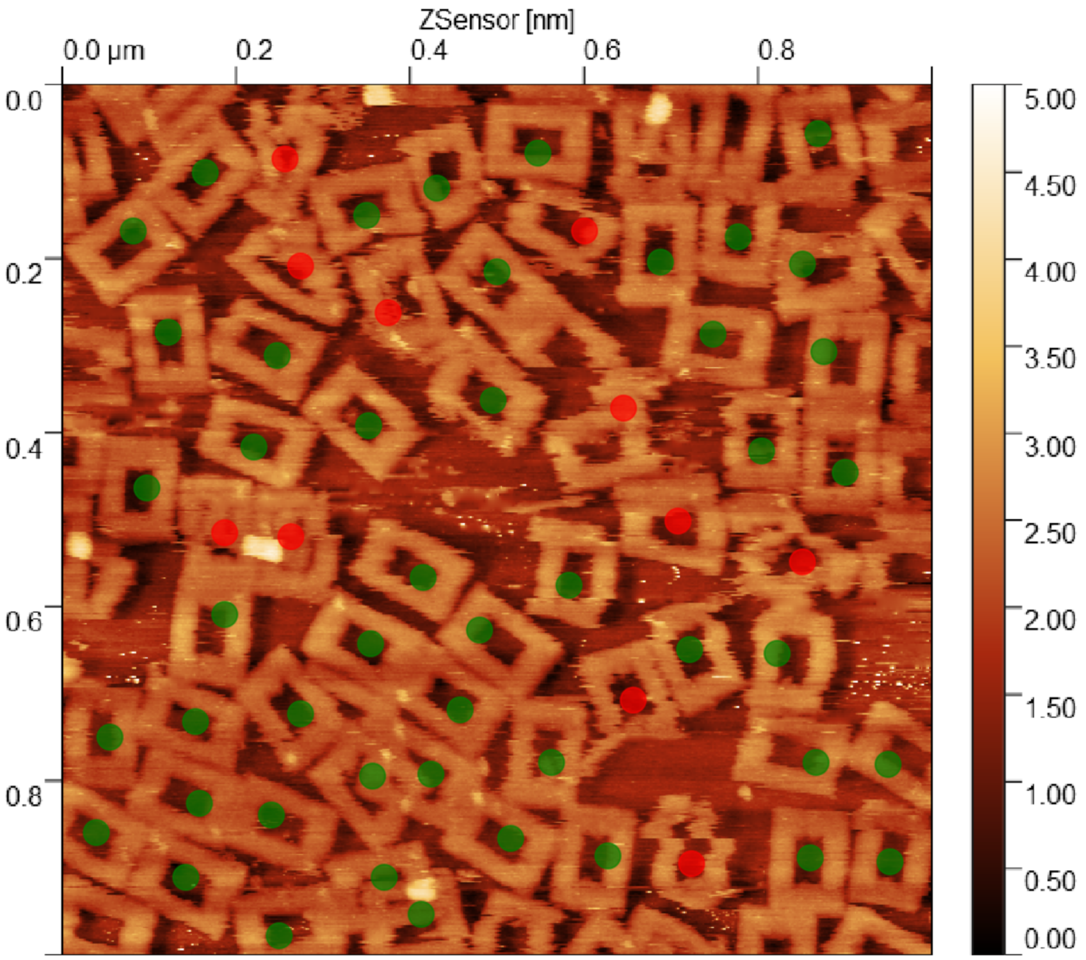

Figure S21: Frame purified Fraction 3 (300µL EV)5x5µm AFM image
